# GAN-MAT: Generative Adversarial Network-based Microstructural Profile Covariance Analysis Toolbox

**DOI:** 10.1101/2023.04.20.537642

**Authors:** Yeong Jun Park, Mi Ji Lee, Seulki Yoo, Chae Yeon Kim, Jong Young Namgung, Yunseo Park, Hyunjin Park, Eun-Chong Lee, Yeo Dong Yun, Casey Paquola, Boris C. Bernhardt, Bo-yong Park

**Author notes:** **Corresponding Author:** Bo-yong Park, PhD, Department of Data Science, Inha University, Incheon, Republic of Korea.

## Abstract

Multimodal magnetic resonance imaging (MRI) provides complementary information for investigating brain structure and function; for example, an *in vivo* microstructure-sensitive proxy can be estimated using the ratio between T1- and T2-weighted structural MRI. However, acquiring multiple imaging modalities is challenging in patients with inattentive disorders. In this study, we proposed a comprehensive framework to provide multiple imaging features related to the brain microstructure using only T1-weighted MRI. Our toolbox consists of (i) synthesizing T2-weighted MRI from T1-weighted MRI using a conditional generative adversarial network; (ii) estimating microstructural features, including intracortical covariance and moment features of cortical layer-wise microstructural profiles; and (iii) generating a microstructural gradient, which is a low-dimensional representation of the intracortical microstructure profile. We trained and tested our toolbox using T1- and T2-weighted MRI scans of 1,104 healthy young adults obtained from the Human Connectome Project database. We found that the synthesized T2-weighted MRI was very similar to the actual image and that the synthesized data successfully reproduced the microstructural features. The toolbox was validated using an independent dataset containing healthy controls and patients with episodic migraine as well as the atypical developmental condition of autism spectrum disorder. Our toolbox may provide a new paradigm for analyzing multimodal structural MRI in the neuroscience community, and is openly accessible at https://github.com/CAMIN-neuro/GAN-MAT.

## Introduction

Multimodal magnetic resonance imaging (MRI) enables the investigation of brain structure and function, and their relationships, *in vivo*. Using structural MRI of T1-weighted (T1w) and T2-weighted (T2w) images, we can assess the anatomical features of the brain, such as cortical thickness, curvature, and volume. Both T1w and T2w data elucidate brain structures, but the image contrast is a major difference. In T1w MRI, the white matter is bright and the gray matter is dark, whereas T2w imaging shows the opposite intensity patterns. This is a consequence of the different MRI parameters. For example, in T1w, the repetition time (TR), which is the time between successive pulse sequences applied to the same slice, and the echo time (TE), which is the time from the center of the radio-frequency pulse to the center of the echo, were shorter than those in T2w MRI. A short TR leads to a strong T1 weighting, whereas a long TE results in a strong T2 weighting. Indeed, T1 and T2 properties are heterogeneous across different tissue types according to the amount of available free water, yielding different image contrasts (Stanisz et al., 2005).

In addition to brain morphology, we can assess microstructural information of the brain using T1w and T2w MRI without obtaining cytoarchitecture data. Specifically, an approximation of the brain microstructure can be estimated using a microstructure-sensitive proxy, which is calculated based on the ratio between T1w and T2w imaging contrasts (Glasser et al., 2014; Glasser and van Essen, 2011), thus enabling us to investigate the cortical microstructure *in vivo*. A recent study suggested a method for analyzing the interregional relations of the brain microstructure (Paquola et al., 2019) by calculating cortical layer-wise microstructural profiles among different brain regions. By applying dimensionality reduction techniques, they generated a low-dimensional representation of the cortical microstructure, referred to as a microstructural gradient. The microstructural gradient represents a well-known hierarchical cortical model of the sensory-fugal axis, which expands from the sensory regions to the limbic cortices (Mesulam, 1998; Paquola et al., 2019). This feature has been widely adopted to assess the microstructural profiles of the brain in healthy adults as well as during adolescence (Paquola et al., 2019, n.d.; Whitaker et al., 2016). Indeed, the microstructural gradient linked the macroscale connectome to microscale cell-type-specific expression during adolescent development, suggesting the validity of this feature for investigating multiscale properties of the brain (Paquola et al., n.d.). However, it is difficult to obtain microstructural features because it requires the acquisition of both T1w and T2w MRI, which is time-consuming and costly. Moreover, obtaining multiple imaging data within a restricted time may not be possible for individuals with psychiatric disorders showing inattentive behaviors, such as autism spectrum disorder and attention-deficit/hyperactivity disorder. Because of these issues, many open databases and clinics typically provide only T1w MRI, not T2w, for research (di Martino et al., 2017, 2014; Milham et al., 2012; Nooner et al., 2012). One approach for mitigating this limitation is image synthesis. If we can synthesize T2w MRI images from T1w images, we can generate a microstructural gradient using only T1w MRI images with reduced time and cost.

Image synthesis has been conducted in many prior works using natural images and texts (Huang et al., 2018; Sangkloy et al., n.d.; Thies et al., 2019; Wang et al., n.d.; Zhang et al., 2019), and transferred to study medical imaging data (Chira et al., 2022; Nie et al., 2018; Osokin et al., 2017; Shin et al., 2018). For example, one study generated high-resolution images from low-resolution data using deep variational autoencoders (Chira et al., 2022), and another work generated brain MRI with tumors using a generative adversarial network (GAN) (Huang et al., n.d.; Osokin et al., 2017). Additionally, one study synthesized multiple imaging modalities using GAN, such as computed tomography from MRI, 7T MRI from 3T MRI, and T2w from T1w MRI (Nie et al., 2018). GAN is a deep learning model synthesizing new imaging data consisting of a generator and discriminator (Goodfellow et al., 2014). The GAN model generates data by simultaneously training both the generator and discriminator. The generator makes fake images as similar as possible to the original image, and the discriminator distinguishes whether the input images are fake or real. The generator and discriminator are adversarial. Specifically, the generator is trained to make the discriminator fail to classify between fake and real data, and the discriminator is trained to distinguish between real and fake images as accurately as possible. A recent study introduced a conditional GAN by adding specific conditions to the vanilla GAN (Nie et al., 2018). One representative model of the conditional GAN is pix2pix, which processes paired data of input and label images (Isola et al., 2016), and another model, called CycleGAN, allows the unpaired set of images (Zhu et al., 2017).

Several studies have proposed models for synthesizing T2w MRI images from T1w MRI images(Kawahara and Nagata, 2021; Yang et al., 2020; Zhao et al., 2021). However, these studies are limited to yielding two-dimensional (2D) MRI data and focus on improving the accuracy of image synthesis without providing a comprehensive framework to study the brain microstructure *in vivo*. In this study, we propose a toolbox consolidating (i) the synthesis of T2w MRI images from T1w images using a conditional GAN, (ii) the calculation of a microstructure-sensitive proxy, and (iii) the generation of ready-to-use microstructural features (**Fig. 1A**).

**Fig. 1.**
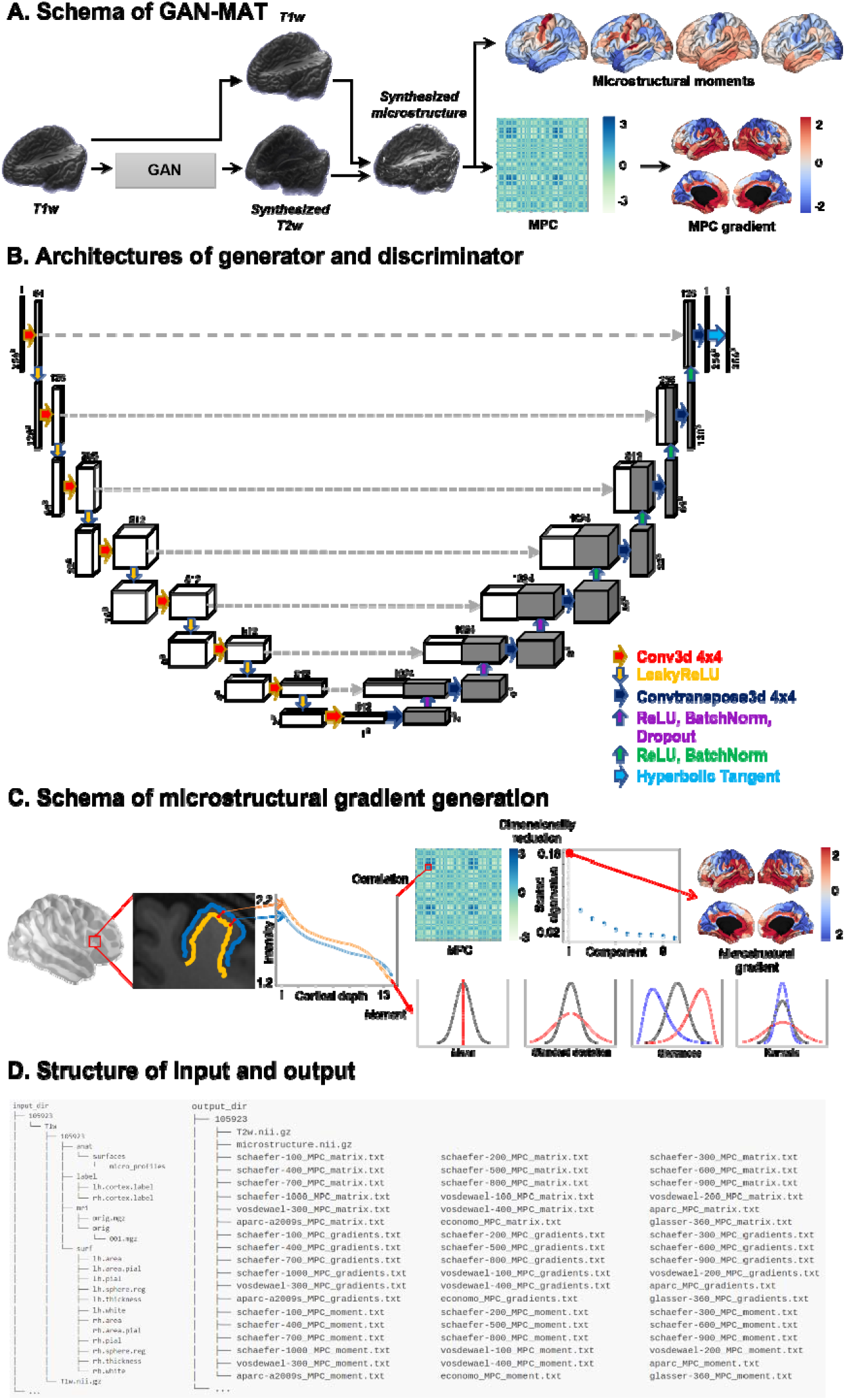
A proposed toolbox synthesizing T2w MRI from T1w MRI and estimating brain microstructural features. **(A)** Shown is the schema of our toolbox, the Generative Adversarial Network-based Microstructural Profile Covariance Analysis Toolbox (GAN-MAT). It contains a conditional GAN synthesizing T2w from T1w MRI and calculates the ratio between T1w and T2w intensities to obtain the microstructure-sensitive proxy. The moment features are calculated from the synthesized microstructure data. The microstructural profile covariance (MPC) matrix is constructed based on the linear correlations of intracortical T1w/T2w intensity profiles between different brain regions, and microstructural gradients are generated using nonlinear dimensionality reduction techniques. **(B)** The architectures of the generator (top) and discriminator (bottom) are shown. **(C)** We describe the schema for generating the microstructural gradient. After generating 14 equivolumetric surfaces (left), we calculate linear correlations of cortical depth-dependent T1w/T2w intensity profiles between different cortical regions to make an MPC matrix (top middle). By applying dimensionality reduction techniques, we generate the microstructural gradient (top right). Additionally, four moment features are calculated from the intracortical microstructural profiles (bottom right). **(D)** Shown is the organization of the input (left) and output (right) directories of a sample subject. *Abbreviations*: T1w, T1-weighted; T2w, T2-weighted; GAN, generative adversarial network; MRI, magnetic resonance imaging.

## Methods

### Imaging data

We studied structural MRI data from three independent sites: (i) Human Connectome Project (HCP) (http://www.humanconnectome.org/) (van Essen et al., 2013), (ii) Samsung Medical Center (SMC), and (iii) Autism Brain Imaging Data Exchange II initiative (ABIDE-II; https://fcon_1000.projects.nitrc.org/indi/abide) (di Martino et al., 2017). The GAN model was constructed using healthy young adults from the HCP dataset and its generalizability was validated using the healthy and diseased populations from the SMC dataset. Finally, we applied the model to the ABIDE-II dataset to confirm its generalizability to both typical and atypical neurodevelopmental conditions. The detailed image acquisition parameters are as follows:

i. HCP: We obtained T1w and T2w data of 1,104 healthy young adults (mean ± standard deviation (SD) age = 28.8 ± 3.7 years; 54.9% female) from the HCP database. The T1w MRI was performed using a 3D magnetization-prepared rapid acquisition gradient echo (MPRAGE) sequence (TR = 2,400 ms; TE = 2.14 ms; field of view (FOV) = 224 × 224 mm^2^; voxel size = 0.7 mm isotropic; number of slices = 256), and the T2w MRI was performed using a 3D T2-SPACE sequence (TR = 3,200 ms; TE = 565 ms; FOV = 224 × 224 mm^2^, voxel size = 0.7 mm isotropic, number of slices = 256).
ii. SMC: From the SMC site, we obtained T1w and T2w data of 43 healthy controls (mean ± SD age = 35.1 ± 7.6 years; 76.7% female) and 58 individuals with migraine (mean ± SD age = 34.3 ± 8.2 years; 75.8% female). The T1w MRI was scanned turbo field echo (TFE) sequence (TR = 8.2 ms; TE = 3.8 ms; field of view (FOV) = 256 × 256 mm^2^; voxel size = 1.0 mm isotropic; and the number of slices = 180), and the T2w MRI was scanned turbo spin echo (TSE) sequence (TR = 3,000 ms; TE = 280 ms; FOV = 256 × 256 mm^2^, voxel size = 1.0 mm isotropic, and the number of slices = 180).
iii. ABIDE-II: Additionally, we obtained T1w data of 36 neurotypical controls (mean ± SD age = 13.0 ± 4.6 years; 2.8% female) and 47 individuals with autism (mean ± SD age = 11.46 ± 5.7 years; 10.6% female) from the ABIDE-II database, which was the same list as our previous study (Park et al., 2021a). As the database did not provide T2w MRI data, we only studied T1w data. Study participants were recruited from two independent sites: New York University Langone Medical Center (NYU) and Trinity College, Dublin (TCD). The T1w MRI was scanned using 3D MPRAGE sequence for both NYU (TR = 2,530 ms; TE = 3.25 ms; FOV = 256 × 256 mm^2^; voxel size = 1.3 × 1.0 × 1.3 mm^3^; and number of slices = 128) and TCD (TR = 8,400 ms; TE = 3.90 ms; FOV = 230 × 230 mm^2^; voxel size = 0.9 mm isotropic; and number of slices = 180).

### MRI data preprocessing

i. HCP: The HCP data were preprocessed using the minimal preprocessing pipelines for HCP (Glasser et al., 2013). The T1w and T2w data were corrected for gradient nonlinearity and b0 distortions and co-registered using a rigid-body transformation. Bias field correction was performed based on the inverse intensities from T1- and T2-weighting. The processed data were nonlinearly registered to the standard Montreal Neurological Institute (MNI152) space. White and pial surfaces were generated by following the boundaries between the different tissues (Dale et al., 1999; Fischl et al., 1999b, 1999a). The mid-thickness surface was generated by averaging the white and pial surfaces, and was used to generate the inflated surface. The spherical surface was registered to Conte69 template using MSMAll (Glasser et al., 2016).
ii. SMC: The SMC data were preprocessed using the Fusion of Neuroimaging Preprocessing (FuNP) pipeline (Park et al., 2019) that integrated AFNI, FSL, FreeSurfer, and ANTs (Avants et al., 2011; Cox, 1996; Fischl, 2012; Glasser et al., 2013; Jenkinson et al., 2012). Similar to the HCP pipeline, gradient nonlinearity and b0 distortion correction, nonbrain tissue removal, and intensity normalization were performed. White matter, pial, and mid-thickness surfaces were generated, and the inflated surface was spherically registered to the Conte69 template with 164k vertices and down-sampled to a 32k vertex mesh.
iii. ABIDE-II: T1w data from the ABIDE-II database were preprocessed using FreeSurfer (Dale et al., 1999; Fischl, 2012; Fischl et al., 2001, 1999a, 1999b; Ségonne et al., 2007), which included gradient non-uniformity correction, non-brain tissue removal, intensity normalization, and tissue segmentation. White and pial surfaces were generated, and topology correction, inflation, and spherical registration to the fsaverage template were performed.

### T2w image synthesis using T1w MRI

We constructed a conditional GAN-based model to synthesize T2w MRI images from T1w MR images using the HCP individuals. The participants were randomly divided into training (n = 752), validation (n = 187), and test (n = 165) datasets. The conditional GAN was based on the pix2pix, which was developed for 2D data (Isola et al., 2016), and its architecture was adjusted to process 3D input data (**Fig. 1B**). The generator was based on the U-Net architecture, which has skip connections linking the layers between the encoder and decoder (Isola et al., 2016). First, the original 3D T1w data were registered onto a 0.7 mm isotropic MNI152 standard space with a matrix size of 260 × 311 × 260 using GNU parallel (TANGE, 2018) and resized to 256 × 256 × 256. The data were processed using seven 3D convolutional layers with a leaky rectified linear unit (LeakyReLU) activation function. At the end of the encoding phase, the latent feature was convoluted and 512 units were obtained. Three deconvolutions with RELU, batch normalization, and dropout with a ratio of 0.5 were applied, and four additional deconvolutions with RELU and batch normalization were conducted. Finally, deconvolution and a hyperbolic tangent were applied, and 256 × 256 × 256 output data were generated. The skip connection was linked to the corresponding decoding layer at each encoding layer. The discriminator was constructed using PatchGAN (Isola et al., 2016). It discriminates images in units of patches; thus, it is faster than conventional discriminators that distinguish entire images simultaneously. The 256 × 256 × 256 input matrix was passed through five convolution layers, and finally, a sigmoid function was applied. To optimize the hyperparameters, we adopted the Adam optimizer, which is a stochastic gradient descent method. The objective function was defined as follows:

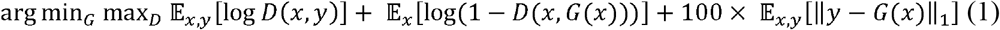

where *G* and *D* denote the generator and discriminator and *x* and *y* denote the input and label images, respectively. The objective function comprises the following three terms: the first determines whether an input is fake or real. The second term is an adversarial term in which the discriminator receives a fake image generated from the generator. The generator is trained to create fake images to enable the discriminator to classify the fake image as real, and the discriminator is trained to accurately distinguish between fake and real images. The last term is the regularization term for training stability. We assessed the performance of the model by calculating the mean absolute error (MAE) between the actual and synthesized T2w data after normalizing the image intensities between 0 and 1. In addition to the global performance, we assessed the tissue type-specific synthesis performance by calculating the MAE of gray matter, white matter, and cerebrospinal fluid. Regional performance was assessed in the frontal, temporal, parietal, occipital, insular, and limbic cortices, and subcortical structures were defined using the Brainnetome atlas (Fan et al., 2016).

### Microstructural profile of the brain

We generated a microstructure-sensitive proxy based on the ratio between the actual T1w and synthesized T2w contrasts (Glasser et al., 2014; Glasser and van Essen, 2011) (**Fig. 1C**). To calculate the intracortical microstructure profiles, we first generated 14 equivolumetric surfaces within the cortex between the inner white and outer pial surfaces, and then sampled the T1w/T2w intensities along these surfaces(Paquola et al., 2019). The statistical moment features (mean, SD, skewness, and kurtosis) were then calculated from the intensity profiles (Paquola et al., n.d.; Schleicher et al., 2009). The mean and SD indicate the overall distribution of the intracortical microstructure, skewness represents shifts in intensity values toward the supragranular layers (i.e., positive) or flat distribution (i.e., negative), and kurtosis indicates whether the tails of the intensity distribution contain extreme values. In addition to the moment features, we constructed a microstructural profile covariance (MPC) matrix by calculating the linear correlations of cortical depth-dependent T1w/T2w intensity profiles between different cortical regions defined using the Schaefer atlas with 300 parcels (Schaefer et al., 2018), while controlling for the average whole-cortex intensity profile(Paquola et al., 2019). The matrix was thresholded to zero and log-transformed (Paquola et al., 2019). We generated a microstructural gradient from the MPC matrix, which is a low-dimensional representation of connectome organization that explains the spatial variation in connectivity (Margulies et al., 2016; Paquola et al., 2019)using the BrainSpace toolbox (https://github.com/MICA-MNI/BrainSpace) (Vos de Wael et al., 2020). Specifically, we applied diffusion map embedding after applying a normalized angle kernel to the group-averaged MPC matrix, leaving the top 10% of elements for each brain region [48]. Diffusion map embedding is a nonlinear dimensionality reduction technique that is robust to noise and computationally efficient compared with other nonlinear manifold learning techniques(Tenenbaum et al., 1995; von Luxburg, 2007). It is controlled by two parameters, α and *t*, where α controls the influence of the density of the sampling points on the manifold (α = 0, maximal influence; α = 1, no influence), and *t* scales the eigenvalues of the diffusion operator. The parameters were set as α = 0.5 and *t* = 0 to retain the global relations between data points in the embedded space, following prior applications (Hong et al., 2019; Margulies et al., 2016; Paquola et al., 2019; Park et al., 2021b; Vos de Wael et al., 2020). Individual gradients were estimated and aligned to the template gradient using Procrustes alignment (Langs et al., 2015; Vos de Wael et al., 2020). We evaluated the similarity between the actual and synthesized microstructural moments and gradient features based on linear correlations, where the significance of the correlation was assessed using 1,000 spin permutation tests that accounted for spatial autocorrelation (Alexander-Bloch et al., 2018; Vos de Wael et al., 2020). Additionally, we assessed the global MAE between the actual and synthesized microstructural features.

### Generalizability of the model using an independent dataset

To assess the reliability and robustness of our toolbox, we applied the HCP-driven model to an independent SMC dataset containing healthy controls and individuals with migraine. We synthesized T2w from T1w and calculated the microstructure-sensitive proxy and relevant moment and gradient features. Performance was evaluated using the MAE between the actual and synthesized T2w images and the linear correlations between the actual and synthesized microstructural features.

### Application to the developmental conditions

We applied the toolbox to T1w MRI of neurotypical controls and individuals with autism obtained from the ABIDE-II database (di Martino et al., 2017) to synthesize T2w data. As the ABIDE-II database did not provide T2w MRI data, we stratified the synthesized microstructural gradient values according to four cortical hierarchical levels (idiotypic, unimodal association, heteromodal association, and paralimbic) (Mesulam, 1998) to assess whether the gradient followed a well-known sensory-fugal brain hierarchy (Paquola et al., 2019).

### Sensitivity analyses

i. *Bootstrap analysis*. We trained the GAN model using different training and validation datasets and synthesized T2w MRI images from the T1w data. We assessed the performance of the model by calculating the MAE between the actual and synthesized T2w images as well as the microstructure-sensitive proxy (T1w/T2w ratio) of the test dataset. The analysis was repeated ten times.
ii. *Two-dimensional model*. In the main analyses, we modified the original 2D-based pix2pix model to process the 3D data. Additionally, we tested whether the original pix2pix model could synthesize T2w images. To this end, we sliced the 3D T1w images along each axis (x, y, and z). The original model consisted of one discriminator; however, we used three discriminators to distinguish the sliced images along each axis. Three synthesized images from the x-, y-, and z-axes were merged in the final stage to yield the 3D data. Model performance was assessed using the MAE between the actual and synthesized T2w and T1w/T2w ratios.
iii. *Synthesis of T1w/T2w ratio*. In addition to synthesizing T2w MRI images from T1w, we trained the GAN to synthesize T1w/T2w directly. We tested both the 3D and 2D models and calculated the MAE to assess the performance.

## Results

### Organization of the toolbox

The developed toolbox requires the input data to be organized in a specific format containing T1w data and several FreeSurfer output files (**Fig. 1D**). The toolbox can be implemented using a single command *“gan-mat -input_dir /INPUT/DATA/DIRECTORY -output_dir/OUTPUT/DIRECTORY”*. It then yields brain microstructural and intracortical moment features as well as the MPC matrix and its microstructural gradient, mapped onto 18 different parcellation schemes (Cruces et al., 2022).

### Synthesis of T2w from T1w MRI

We synthesized 3D T2w MRI images from T1w data using a modified pix2pix model. The trained model was applied to the holdout test dataset and the actual and synthesized T2w images showed similar spatial patterns (mean ± SD MAE of the whole brain across individuals = 0.036 ± 0.010) when the image intensity of each subject was scaled between 0 and 1 (**Fig. 2A**). The synthesis performance was slightly different among the tissue types, with the best performance observed in the white matter (white matter = 0.032 ± 0.009, gray matter = 0.038 ± 0.008, and cerebrospinal fluid = 0.041 ± 0.005 across individuals). When we stratified the MAE according to different lobes and subcortical structures, the subcortical structures showed the best performance, whereas the parietal and occipital lobes showed relatively higher errors but were still comparable.

**Fig. 2.**
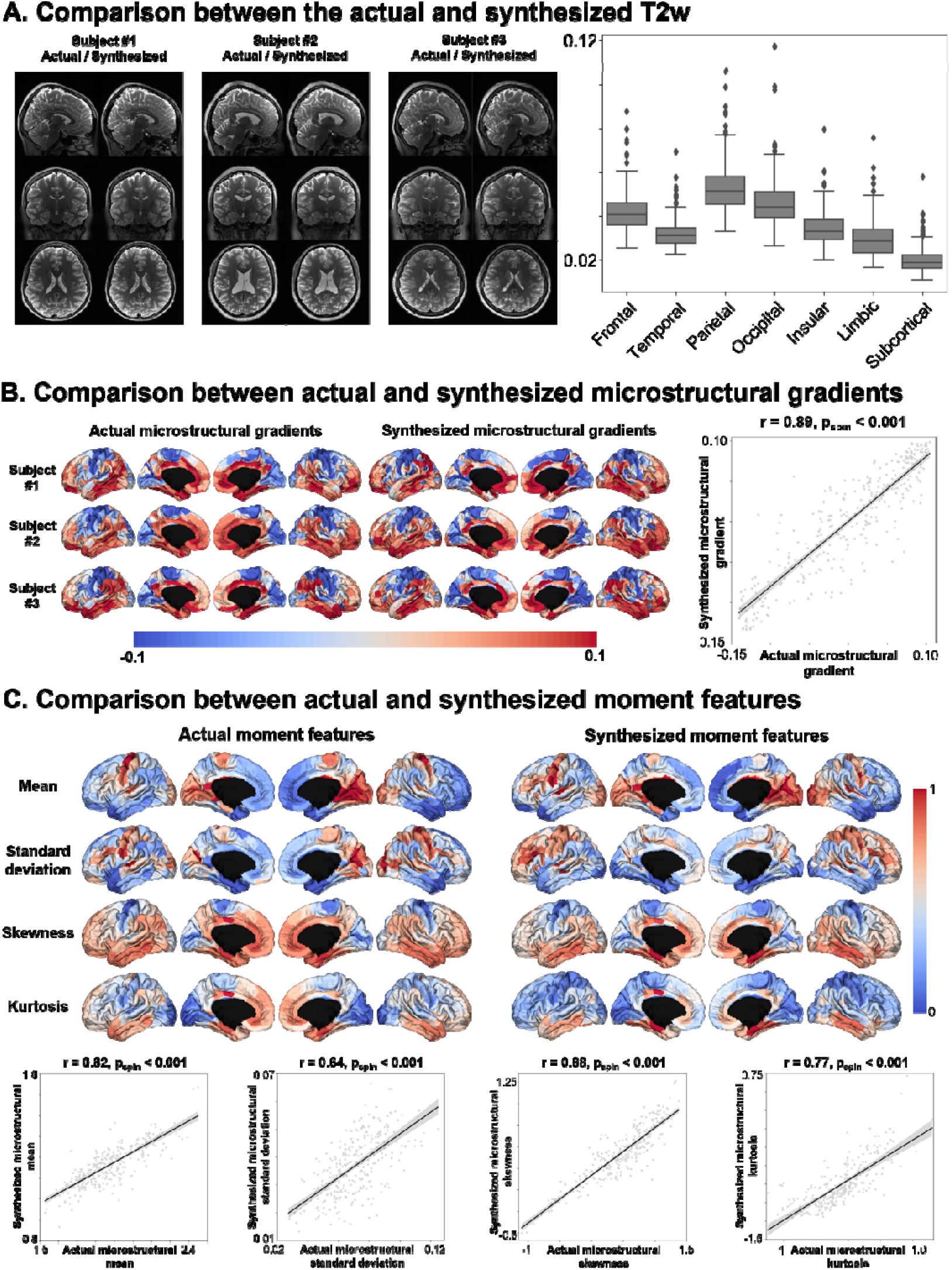
Performance of the synthesis model. **(A)** Visualization of the actual and synthesized T2w images of three representative participants (left). The box plots show the MAE of different cortical and subcortical structures (right). **(B)** The actual (left) and synthesized (middle) microstructural gradients of three representative participants are shown. The similarity of the group-level gradients is assessed using spatial correlations with spin permutation tests (right), where the gray area indicates a 95% confidence interval. **(C)** The actual (left) and synthesized (right) group-level moment features are shown. The group-level correlations between actual and synthesized moment features are shown (bottom). *Abbreviations*: MAE, mean absolute error; T2w, T2-weighted.

### Synthesized brain microstructure

The validity of the synthesized T2w images was evaluated by assessing the similarity between the actual and synthesized microstructure-sensitive proxies based on the T1w/T2w ratio (**Fig. 2B**). We observed a high similarity in the synthesized microstructure-sensitive proxy, where the mean ± SD MAE was 0.007 ± 0.002 across individuals. Moreover, the generated microstructural gradient showed a well-known sensory-fugal hierarchy that radiated from sensory and motor areas with higher myelination toward heteromodal associations and paralimbic regions with lower myelin content. The individual-level correlations between the actual and synthesized microstructural gradients showed significant associations (mean ± SD correlation coefficient = 0.71 ± 0.05, p_spin_ < 0.001). Additionally, we assessed the similarity between the actual and synthesized moment features, which also showed high similarities (mean: mean ± SD correlation coefficient = 0.70 ± 0.07, p_spin_ < 0.001; SD: 0.53 ± 0.08, p_spin_ < 0.001; skewness: 0.77 ± 0.06, p_spin_ < 0.001; kurtosis: 0.63 ± 0.12, p_spin_ < 0.001; **Fig. 2C**).

### Validation of the model using an independent dataset

The generalizability of the model was evaluated by applying it to an independent SMC dataset. The MAE between the actual and synthesized T2w images showed comparable results (mean ± SD MAE across healthy controls = 0.086 ± 0.016; individuals with migraine = 0.089 ± 0.014; **Fig. 3A**). The linear correlations were also comparable between the actual and synthesized microstructural gradients (healthy controls = 0.74 ± 0.04, individuals with migraine = 0.75 ± 0.06) and moment features (healthy controls/individuals with migraine: mean = 0.08 ± 0.04/0.09 ± 0.07, SD = 0.45 ± 0.14/0.47 ± 0.15, skewness = 0.72 ± 0.06/0.75 ± 0.05, kurtosis = 0.60 ± 0.09/0.64 ± 0.10) except for the mean moment feature (**Fig. 3B-C**). These results indicated that our toolbox can be used to investigate the microstructural profiles of both healthy controls and patients with neurological conditions.

**Fig. 3.**
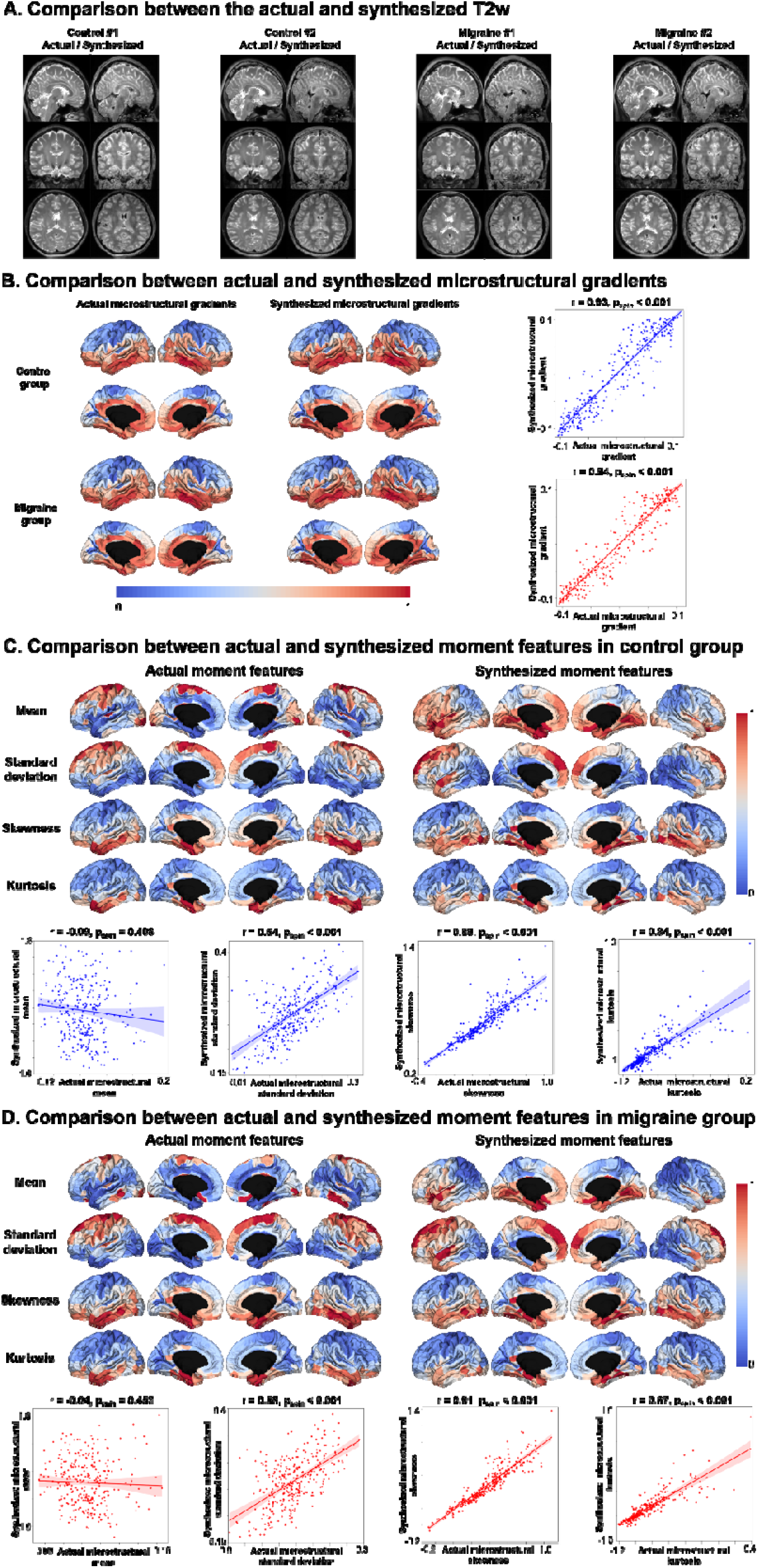
Validation of the toolbox using an independent dataset. **(A)** Visualization of the actual and synthesized T2w images of two representative participants in each group. **(B)** Shown are the microstructural gradients of the control and migraine groups. The group-level correlations between the actual and synthesized gradients are shown with scatter plots. **(C)** We described moment features of the control and **(D)** migraine groups, where the group-level correlations are reported with scatter plots. *Abbreviations*: T2w, T2-weighted.

### Application of the model to the typical and atypical developmental conditions

We applied our toolbox to data from neurotypical controls and individuals with autism which we obtained from the ABIDE II database (di Martino et al., 2017). We estimated the MPC matrix and microstructural gradients for each subject and averaged them to obtain group-representative data for the control and autism groups (**Fig. 4A**). The generated microstructural gradients exhibited a sensory-fugal axis in both groups. When we stratified the gradient values according to the four cortical hierarchical levels (Mesulam, 1998), a clear hierarchy along the cortex was observed, expanding from the lower-level idiotypic to the higher-order association and paralimbic areas (**Fig. 4B**). Together, these results indicate that our toolbox can be generalized to independent datasets of typical and atypical developmental conditions.

**Fig. 4.**
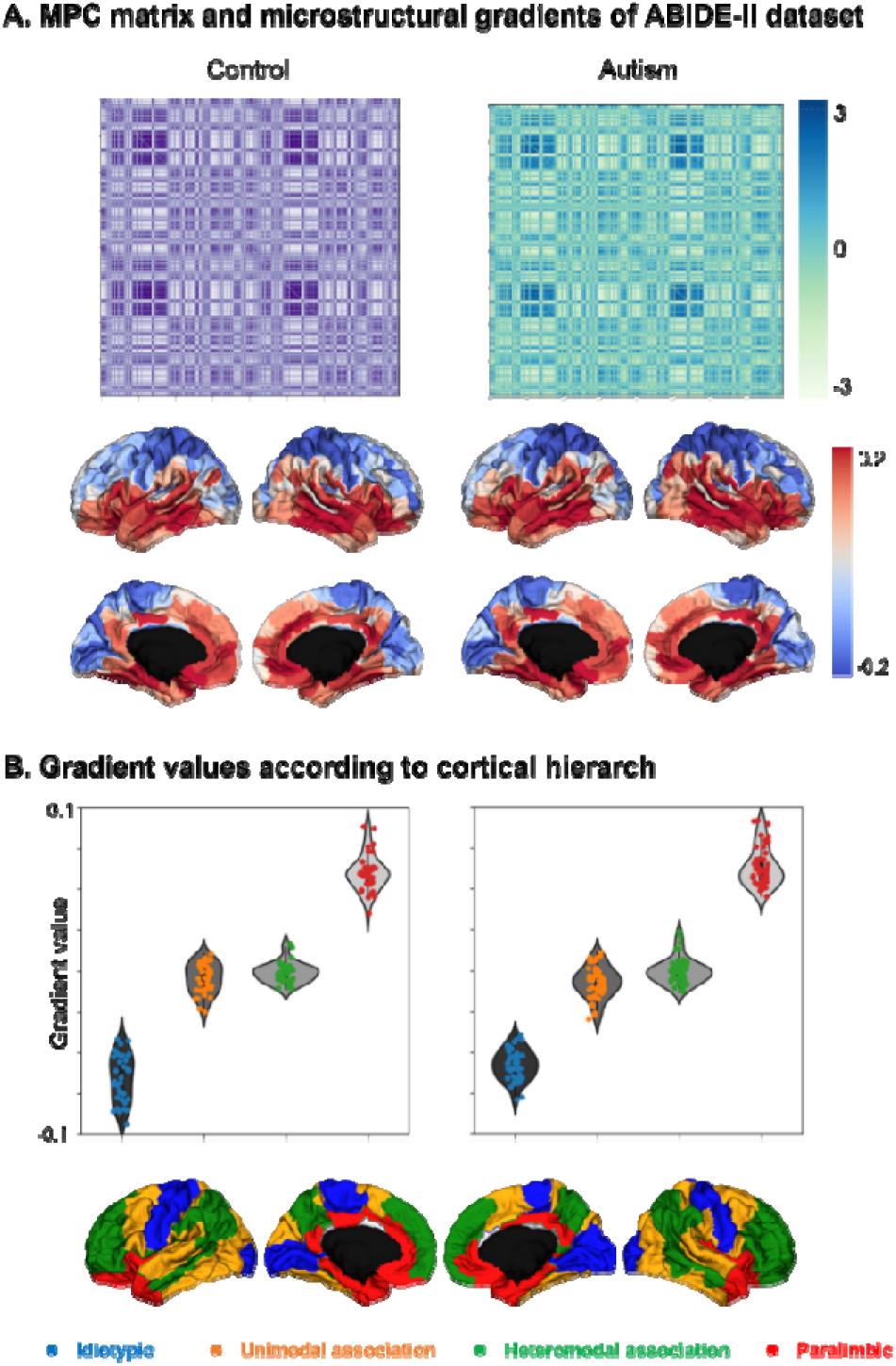
Brain microstructure of an independent dataset. **(A)** We constructed a microstructural profile covariance (MPC) matrix (top) and microstructural gradient (bottom) of neurotypical controls and individuals with autism. **(B)** We stratified the gradient values according to four cortical hierarchical levels.

### Sensitivity analyses

i. *Bootstrap analysis*. We synthesized T2w data from T1w MRI images using randomly selected training and validation datasets to avoid subject-selection bias. The mean ± SD MAE between the actual and synthesized T2w of the test datasets across ten bootstraps was 0.045 ± 0.005 and that of the T1w/T2w ratio was 0.016 ± 0.0004, indicating robustness (**Supplementary Fig. 1**).
ii. *Two-dimensional model*. The original 2D pix2pix model was trained instead of the modified 3D model. The mean ± SD MAE between the actual and synthesized T2w was 0.110 ± 0.023, and the T1w/T2w ratio was 0.027 ± 0.023 (**Supplementary Fig. 2A**). The results based on the 2D model exhibited higher errors than those based on the 3D model.
iii. *Synthesis of T1w/T2w ratio*. Instead of synthesizing T2w, we directly determined the T1w/T2w ratio from the T1w data. The 2D model showed mean ± SD MAE of 0.048 ± 0.008 (**Supplementary Fig. 2B**), and the 3D model showed 0.018 ± 0.002 (**Supplementary Fig. 2C**). These results suggest that synthesizing T2w data is better than synthesizing the T1w/T2w ratio.

## Discussion

The image-synthesis approach benefits neuroimaging studies by generating multiple imaging modalities from a single modal image with reduced time and cost. In this study, we constructed and disseminated a toolbox to analyze the brain microstructure *in vivo* using only T1w MRI. Specifically, the toolbox synthesizes T2w from T1w MRI images and calculates a microstructure-sensitive proxy to generate the MPC matrix, its gradient, and moment features. We observed a high correspondence between the actual and synthesized features, and multiple sensitivity analyses demonstrated the robustness of the toolbox. Our proposed framework may facilitate multimodal neuroimaging studies, particularly for studying brain microstructures using limited neuroimaging modalities.

The concept of image synthesis was introduced in previous neuroimaging studies. For example, one study used a conditional GAN to synthesize T1w from T2w images and T2w from T1w images based on the original pix2pix model (Kawahara and Nagata, 2021). Another study modified the model to process 3D data, in which each dimension was a sagittal, coronal, or axial slice (Zhao et al., 2021). Additionally, a conditional GAN was adopted to improve the quality of the registration and segmentation of brain images containing tumors (Yang et al., 2020). These studies focused on optimizing the distribution of the synthesized image to make it as similar as possible to an actual image. Thus, the aims of these studies were primarily to improve the synthesis accuracy and optimize the hyperparameters of the model. Contrastingly, our work aimed to provide the microstructural features of the brain that can be used in neuroscience and clinical studies to identify markers of specific psychiatric or neurological conditions. For example, the microstructure-sensitive proxy can be used to investigate alterations in brain network organization of Alzheimer’s disease, schizophrenia, epilepsy, and multiple sclerosis (Bernhardt et al., 2018; Boaventura et al., 2022; Ganzetti et al., 2015; Pelkmans et al., 2019; Yasuno et al., 2017), and we can assess behavioral and cognitive traits during typical and atypical development [53]–[55]. Moreover, microstructural features can be used to investigate multiscale neural organization. The microstructural gradient describes macroscopic connectome organization and is associated with gene expression in brain cells (Paquola et al., n.d.; Royer et al., 2020). In summary, our study impacts clinical neuroscience by providing a consolidated framework for synthesizing T2w images from T1w MRI images and generating ready-to-use brain microstructural features.

We demonstrated the reliability and robustness of our toolbox using multiple scenarios. First, we quantitatively tested four different models: (i) synthesis of T2w using a 3D GAN (ii) 2D GAN, (iii) synthesis of the T1w/T2w ratio using a 3D GAN, and (iv) 2D GAN. We found that the first model (3D–T2w synthesis) exhibited the best performance. The superior performance of the 3D model relative to that of the 2D model may be due to the quantity of information. The 2D model uses information on the brain anatomy of each axis (i.e., sagittal, coronal, and axial) for training; thus, it does not consider the geometric properties across different slices. Additionally, synthesizing T2w images is better than directly creating a T1w/T2w ratio. A previous study suggested that the role of T2w images when calculating microstructure-sensitive proxies is to remove blood vessels and dura from the pial surface and reduce the effects of myelin content on pial surface generation via intensity normalization of gray matter (Glasser et al., 2014). If we directly synthesize the T1w/T2w ratio from the T1w data, the GAN model may not consider these biological properties of T2w images, leading to a minor similarity between the actual and synthesized images. Second, we conducted bootstrap tests with different training and validation datasets and found low errors in the gray and white matter. The boundaries between different tissue types and the skull vary largely across individuals, leading to the optimization of hyperparameters for model training. Further studies are required to minimize such errors and improve the synthesis performance. Third, we tested the generalizability of our toolbox by using an independent dataset containing both healthy and diseased populations. These findings indicate that our toolbox is appropriate for investigating disease-related microstructural alterations in the brain using only T1w MRI.

In this study, we developed an end-to-end toolbox for synthesizing T2w images from T1w images and generating brain microstructural features, including moments, MPC matrix, and microstructural gradients. Multiple sensitivity analyses demonstrated the reliability and robustness of this toolbox. Our model may foster future multimodal MRI studies to investigate brain microstructures.

## Data Availability

The imaging and phenotypic data were provided, in part, by the Human Connectome Project, the WU-Minn Consortium (https://www.humanconnectome.org/), and the Autism Brain Imaging Data Exchange initiative (ABIDE-II; https://fcon_1000.projects.nitrc.org/indi/abide/).

## Code Availability

The codes for the GAN-MAT are available at https://github.com/CAMIN-neuro/GAN-MAT.

## Supplementary information

**Supplementary Fig. 1.**
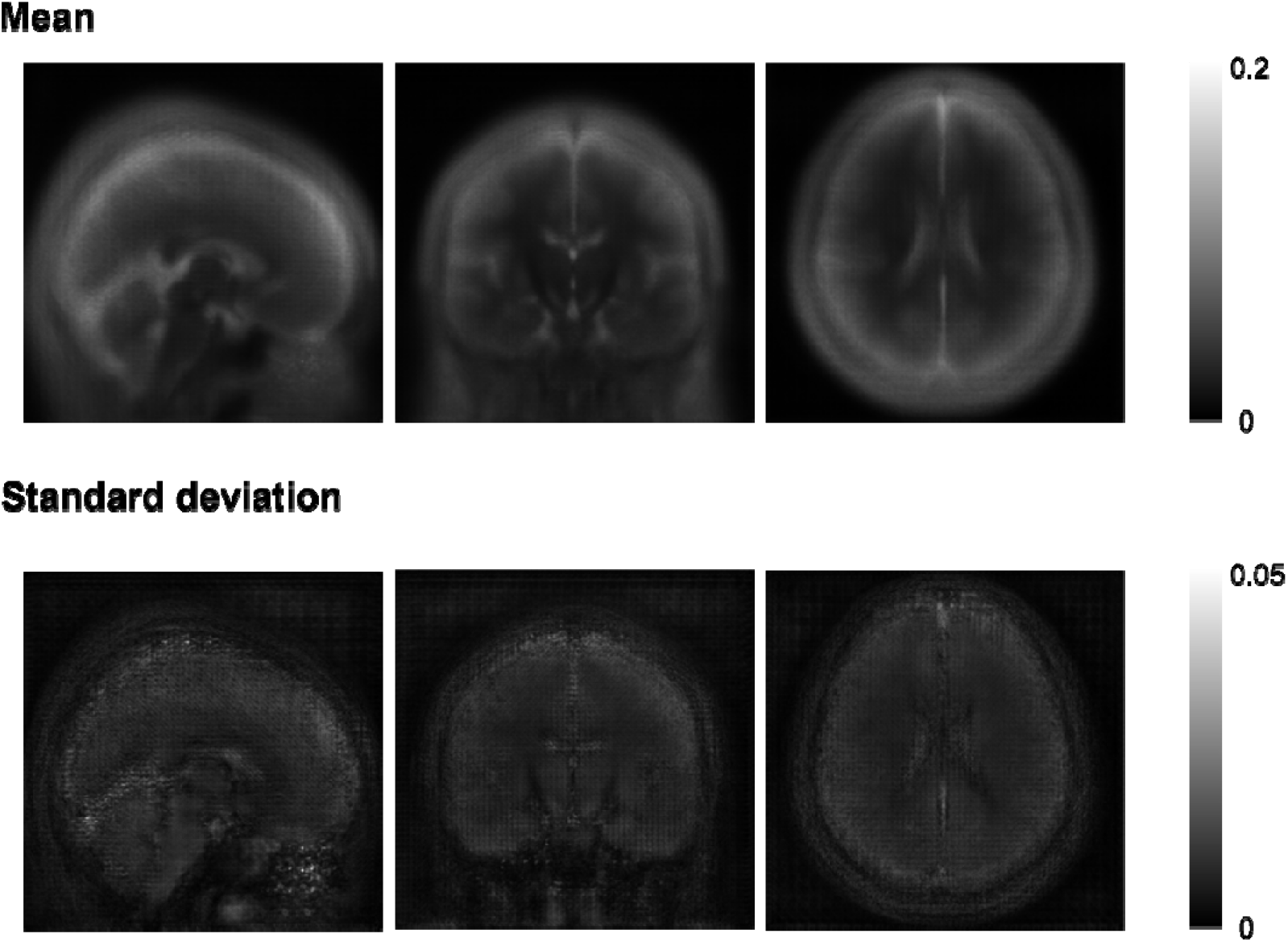
Differences between the actual and synthesized T2-weighted MRI. The mean (top) and standard deviation (bottom) of the differences across ten bootstraps are shown.

**Supplementary Fig. 2.**
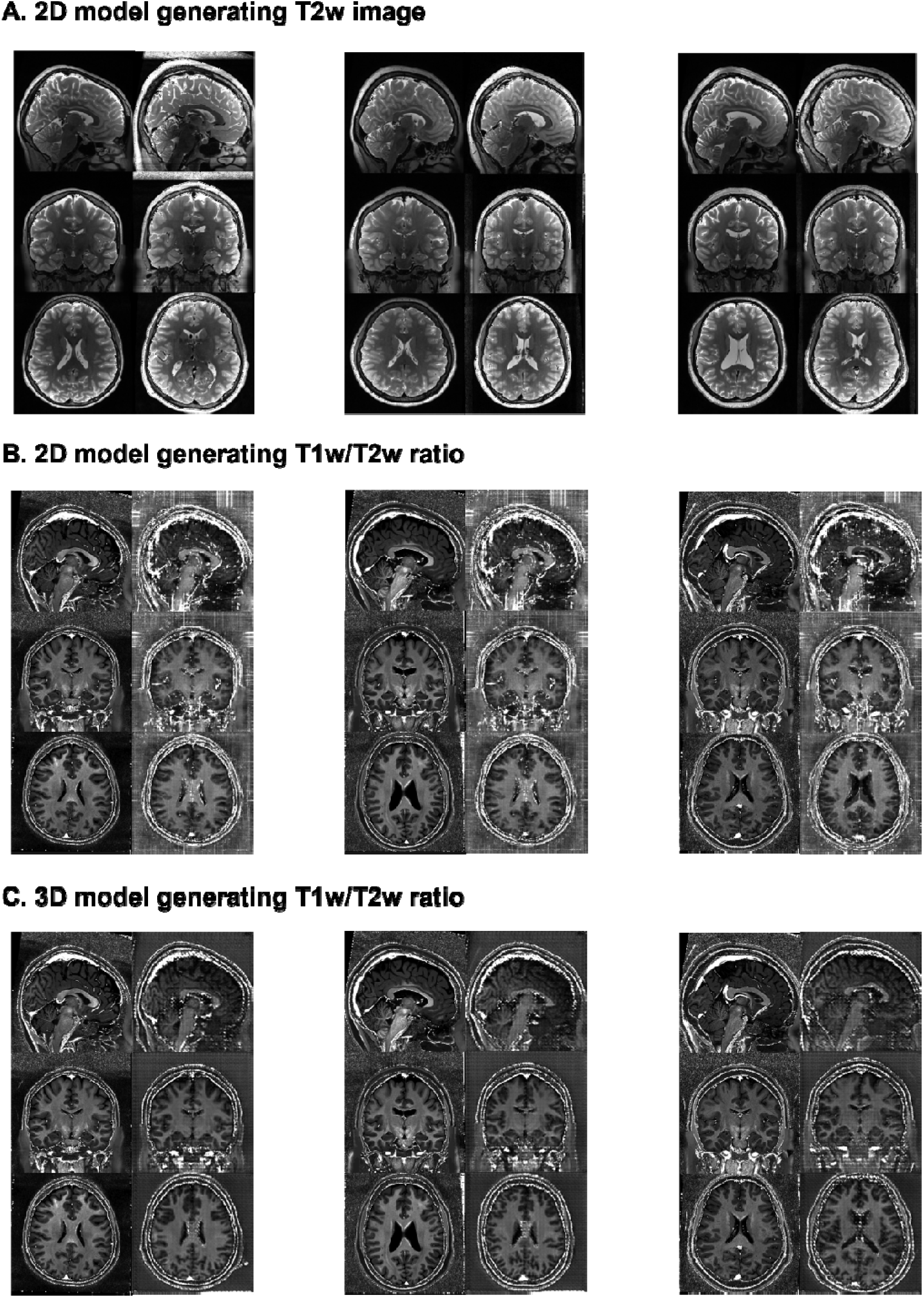
Synthesized images using other models. **(A)** The actual (left), synthesized (right) T2w images using a 2D model, **(B)** T1w/T2w ratio using a 2D model, and **(C)** T1w/T2w ratio using a 3D model of three representative participants. *Abbreviations*: T1w, T1-weighted; T2w, T2-weighted.

